# Metabolic state shapes cortisol reactivity to acute stress: A systematic review and meta-analysis of metabolic and hormonal modulators

**DOI:** 10.1101/2025.08.05.668684

**Authors:** Madeleine Kördel, Maria Meier, Anne Kühnel, Nils B. Kroemer

## Abstract

Individual variability in cortisol stress responses is shaped by multiple physiological factors. Yet the interaction with metabolic and hormonal states remains poorly understood. Therefore, we conducted a systematic review and meta-analysis to examine how metabolic factors (particularly glucose) and sex hormone levels (progesterone and estradiol) influence cortisol reactivity to acute stress. We identified 21 studies (*N* = 1216 participants) and conducted random-effects meta-analyses for metabolic and hormonal states. Across studies, glucose administration was associated with a significant increase in cortisol responses to acute stress compared to fasting or non-glucose control conditions (*d* = 0.30, 95% CI = [0.05, 0.60], BF_10_ = 2.42, NNT = 10.63). In contrast, the effects of sex hormones on cortisol responses were smaller and more variable, with both progesterone and estradiol showing weak and inconsistent associations. Our results highlight a robust modulatory role of metabolic state, specifically glucose availability, on HPA axis reactivity, while evidence for sex hormone effects remains inconclusive. Future research should focus on better harmonization of designs concerning sex hormones and systematically examine interactions between metabolic and hormonal states to better explain sex differences in the prevalences of metabolic and stress-related disorders.

## Introduction

Stress describes a state in which the organism perceives or anticipates physical or psychological threats and initiates an adaptive response (Chrousos, 1992; Goel et al., 2014; Habib et al., 2001; McEwen, 1998). To restore homeostasis, stress systems are activated including the hypothalamic-pituitary-adrenal (HPA) axis and the sympathetic nervous system (Chrousos, 1992; Habib et al., 2001). After stress exposure, the hypothalamus secretes corticotropin-releasing factor, causing the release of adrenocorticotropic hormone from the pituitary, followed by cortisol secretion from the adrenal cortex (Lupien et al., 2005). During acute stress, cortisol helps maintain energy balance by mobilizing glucose to meet immediate energy demands (Sapolsky et al., 2000), and such acute changes can be robustly tracked at the individual level with physiological and neuroimaging measures of stress reactivity (Kühnel et al., 2022). However, chronic or repeated stress can lead to allostatic load, resulting in physiological “wear and tear” (McEwen, 1993). Since activating and regulating stress responses incur additional energetic costs to an organism (Bobba-Alves et al., 2022), acute and chronic stress have been theorized to be modulated by the metabolic state (i.e., glucose metabolism) (Kirschbaum et al., 1997). Similarly, neuroendocrine modulators (e.g., sex hormones) may contribute to individual differences in stress reactivity (Barel et al., 2018; Ulrich-Lai & Herman, 2009) and risk for stress-related mental and metabolic disorders. Although some studies suggest that acute stress can alter insulin sensitivity and increase glucose, estradiol and progesterone levels (Keselman et al., 2017; Kruyt et al., 2012; Lennartsson et al., 2012; Shields et al., 2019), the impact of different metabolic and hormonal states on HPA axis reactivity remains largely elusive, with sparse and inconsistent findings.

To mobilize energy reserves via HPA axis activation during threats, glucose metabolism and stress reactivity are closely intertwined, but questions about the directionality and practical relevance of their interaction remain. On the one hand, it is well established that during acute stress, cortisol secretion raises blood glucose levels to provide energy by stimulating gluconeogenesis and mobilizing carbohydrate stores (Sapolsky et al., 2000). According to the “selfish brain” theory, the brain further reinforces this process by suppressing insulin secretion via cortisol, prioritizing glucose supply to the brain at the expense of peripheral tissues (Peters, 2011; Peters et al., 2004). Consequently, stress-induced increases in glucose metabolism have been demonstrated (Kern et al., 2008). On the other hand, emerging evidence suggests that the current metabolic state, particularly glucose availability, may influence the magnitude of the cortisol response to stress (Sharma et al., 2022; von Dawans et al., 2021; Yajurvedi, 2018). Initial studies have reported strong associations between glucose intake and cortisol responses (Kirschbaum et al., 1997), but more recent studies show smaller or inconsistent effects (Bentele et al., 2021; Gonzalez-Bono et al., 2002; Meier et al., 2021, 2025; von Dawans et al., 2021; Zänkert et al., 2020), while others found no effect (Rüttgens & Wolf, 2022). Glucose administration without a stressor does not increase cortisol levels, suggesting that the effect is specific to stress reactivity rather than a direct action of glucose on cortisol secretion (Kirschbaum et al., 1997). In addition, research on insulin’s role in stress reactivity is scarce and has yielded mixed results so far. While some studies focusing on central effects of insulin report a dampening impact on HPA axis activation (Bohringer et al., 2008), others observed an increased cortisol secretion following peripheral insulin administration (Fruehwald-Schultes et al., 2001). Together, these findings point to a potential effect of metabolic states on the cortisol responses, but a systematic synthesis is lacking to date.

Sex hormones are an additional biological factor potentially modulating stress responses. Throughout the menstrual cycle, hormone levels fluctuate, affecting energy metabolism, leading to interactions with the HPA axis through complex feedback mechanisms that contribute to individual variability and systematic sex differences (Albert & Newhouse, 2019; Barel et al., 2018; Hummel et al., 2023; Kroemer, 2023). For example, low estradiol and progesterone levels in the early follicular phase have been associated with greater subjective distress and slightly elevated basal cortisol (Albert et al., 2015; Klusmann et al., 2022). In this phase, central insulin sensitivity in the hypothalamus is increased (Hummel et al., 2023), potentially fine-tuning stress responses via adapted energy metabolism (Kroemer, 2023). Rising estradiol in the mid-to-late follicular phase appears to be linked to reduced subjective stress reports (Albert et al., 2015). In the luteal phase (characterized by high progesterone), lower basal cortisol but enhanced HPA axis responses to physiological stress have been reported (Klusmann et al., 2022, 2023). Additionally, blood glucose levels are elevated and insulin sensitivity are reduced (Hummel et al., 2023; Lin et al., 2023). Despite some studies suggesting that sex hormones modulate cortisol responses to stress (Barel et al., 2018; Maki et al., 2015; Stephens et al., 2016), findings remain inconsistent (Antov & Stockhorst, 2014; Gordon & Girdler, 2014; Pletzer et al., 2021).

So far, existing meta-analyses on sex hormones have primarily focused on differences in basal cortisol levels (Klusmann et al., 2022) or in response to physiological stressors (Klusmann et al., 2023). However, the effects of metabolic states or sex hormones on cortisol responses to psychosocial stress have not been synthesized so far. To address previous limitations, we conducted a more inclusive meta-analysis that considers the diversity in study designs, hormonal grouping methods, and includes studies regardless of whether an average cortisol response was elicited to avoid systematic biases. To this end, we synthesized studies investigating how metabolic (i.e., glucose and insulin) and sex hormone (i.e., estradiol and progesterone) states influence cortisol responses to acute stressors, including both physical and psychosocial stress induction. Based on previous research, we predicted that glucose administration would enhance cortisol stress responses while sex hormones would modulate these responses in a cycle-dependent manner. Accordingly, we observed a meta-analytic effect of glucose on cortisol responses to stress while modulatory effects of sex hormones did not yield a robust association to date. We argue that clarifying such associations is essential for understanding how metabolic and hormonal states shape HPA axis reactivity to identify promising moderators and mediators of stress-related health outcomes and we derive suggestions for future work to address remaining gaps.

## Methods

The systematic review and meta-analysis were conducted according to the Preferred Reporting Items for Systematic Reviews and Meta-Analyses (PRISMA) 2020 statement (Page et al., 2021). The corresponding preregistration can be found in the International Prospective Register of Systematic Reviews (PROSPERO) database (Registration No. CRD42023465851, https://www.crd.york.ac.uk/prospero/).

### Search strategy

The literature search was conducted in the electronic databases PubMed, Web of Science Core Collection, EBSCO (PsyInfo, PsyINDEX, APA PsycArticles), Livivo, and BIOSIS Citation Index on July 7^th^, 2023. The following search terms were used with a filter for human populations: cortisol AND (stress induction OR acute stress OR stress task OR stress reactivity) AND (progesterone OR estradiol OR insulin OR glucose; for details, see supporting information). The initial database search identified 5478 studies (Figure 1). Three papers were manually added (Bürger et al., 2025; Gonzalez-Bono et al., 2002; Meier et al., 2025), and we selected two eligible reports from a previous meta-analysis (De Souza et al., 1991; Galliven et al., 1997). After removing duplicates and the initial screening of titles, 230 studies remained for further assessment. The screening of abstracts and full-text screening led to the exclusion of 170 studies. We identified 60 eligible studies that fulfilled the inclusion criteria. From these studies, 21 provided sufficient data to be included for the meta-analysis. We used WebPlotDigitizer (Version 5, https://automeris.io/WebPlotDigitizer) to extract data from figures. For studies where sufficient data could not be obtained, we contacted the corresponding authors.

**Fig. 1.**
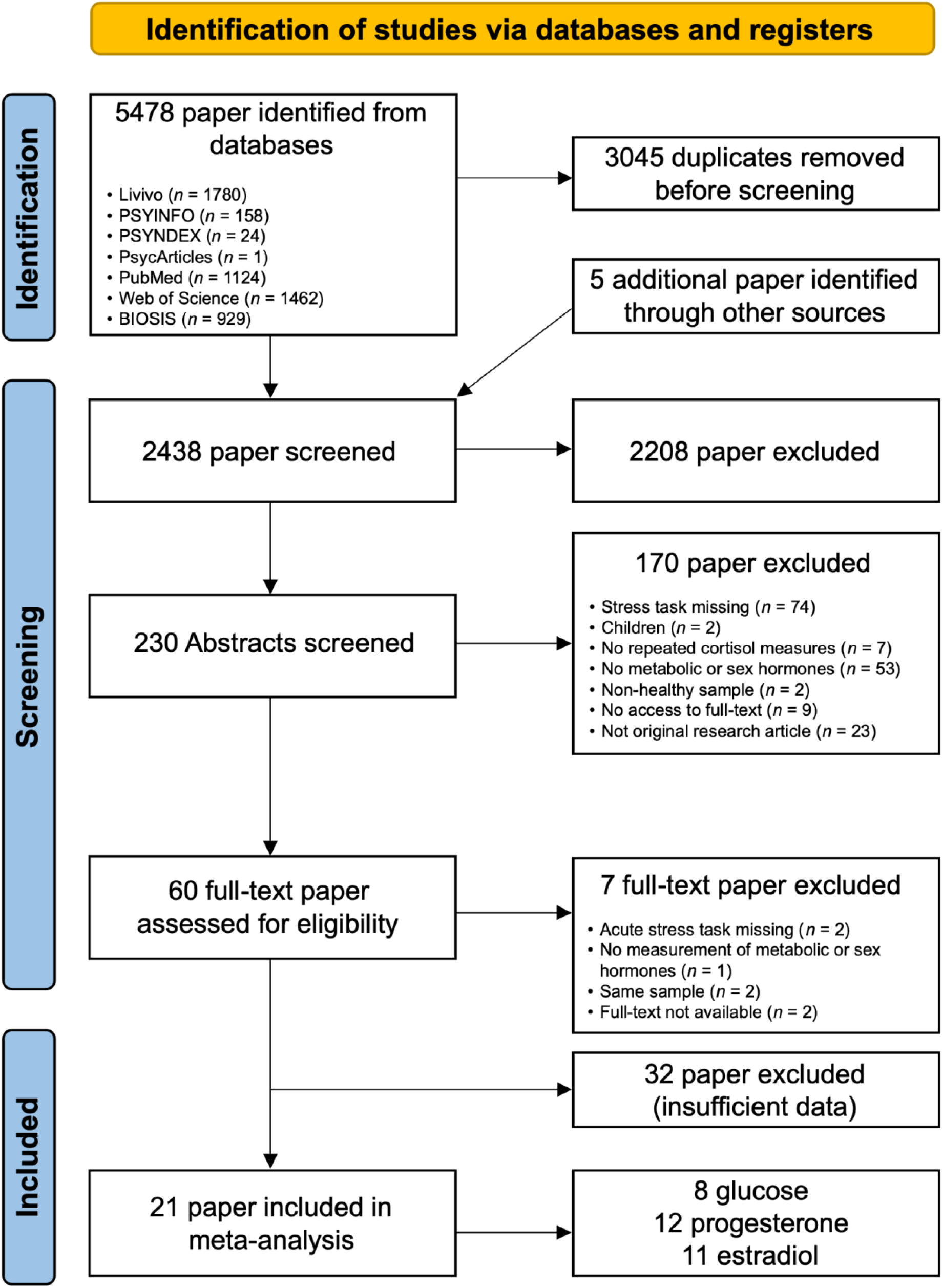
PRISMA flowchart showing the selection criteria for articles screened for inclusion in the meta-analysis.

### Characteristics of metabolic studies

A total of 8 studies (*N* = 458 participants, Table 1) investigated the effects of glucose administration on cortisol stress reactivity. All studies used a between-subjects experimental design to elevate glucose levels with caloric loads (Bentele et al., 2021; Gonzalez-Bono et al., 2002; Kirschbaum et al., 1997; Meier et al., 2021, 2025; Rüttgens & Wolf, 2022; von Dawans et al., 2021; Zänkert et al., 2020). All studies used the Trier Social Stress Test (TSST, Kirschbaum et al., 1993) a standardized psychosocial stress paradigm, except for one, which used a combined psychosocial-physical stressor (Rüttgens & Wolf, 2022), and one used both psychosocial and physical stressors (von Dawans et al., 2021). The majority of experiments were conducted in the afternoon or evening (*n* = 6), with 2 studies scheduled in the morning (Bentele et al., 2021; Meier et al., 2021). Cortisol concentrations were assessed in saliva samples, while capillary blood samples were used to measure glucose levels. In all but one study (Zänkert et al., 2020), the effect of glucose manipulation was confirmed using blood glucose levels. After screening, insulin was excluded from the current meta-analysis due to insufficient data (i.e., one study by Bohringer et al., 2008). Furthermore, no study examined the interaction of sex hormones and glucose on cortisol stress response.

**Table 1.**
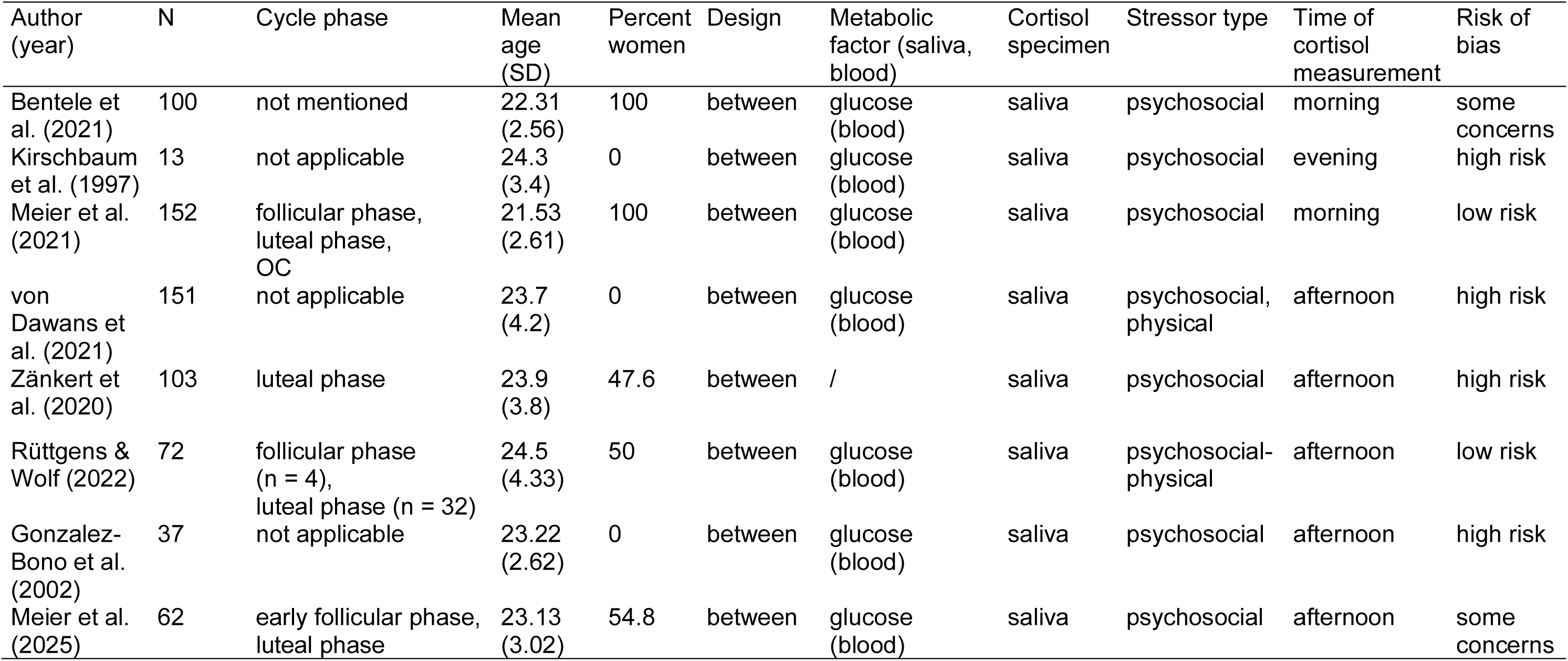
Characteristics of the included studies investigating metabolic states and their effect on cortisol stress reactivity. *Note:* SD = standard deviation. OC = oral contraceptives.

| Author (year) | N | Cycle phase | Mean age (SD) | Percent women | Design | Metabolic factor (saliva, blood) | Cortisol specimen | Stressor type | Time of cortisol measurement | Risk of bias |
| --- | --- | --- | --- | --- | --- | --- | --- | --- | --- | --- |
| Bentele et al. (2021) | 100 | not mentioned | 22.31 (2.56) | 100 | between | glucose (blood) | saliva | psychosocial | morning | some concerns |
| Kirschbaum et al. (1997) | 13 | not applicable | 24.3 (3.4) | 0 | between | glucose (blood) | saliva | psychosocial | evening | high risk |
| Meier et al. (2021) | 152 | follicular phase, luteal phase, OC | 21.53 (2.61) | 100 | between | glucose (blood) | saliva | psychosocial | morning | low risk |
| von Dawans et al. (2021) | 151 | not applicable | 23.7 (4.2) | 0 | between | glucose (blood) | saliva | psychosocial, physical | afternoon | high risk |
| Zänkert et al. (2020) | 103 | luteal phase | 23.9 (3.8) | 47.6 | between | / | saliva | psychosocial | afternoon | high risk |
| Rüttgens & Wolf (2022) | 72 | follicular phase (n = 4), luteal phase (n = 32) | 24.5 (4.33) | 50 | between | glucose (blood) | saliva | psychosocial-physical | afternoon | low risk |
| Gonzalez-Bono et al. (2002) | 37 | not applicable | 23.22 (2.62) | 0 | between | glucose (blood) | saliva | psychosocial | afternoon | high risk |
| Meier et al. (2025) | 62 | early follicular phase, luteal phase | 23.13 (3.02) | 54.8 | between | glucose (blood) | saliva | psychosocial | afternoon | some concerns |

### Characteristics of sex hormone studies

To examine the effects of progesterone and estradiol on cortisol responses to psychological or physical stress, we included studies either comparing women in various hormonal states (e.g., oral contraceptives vs. no oral contraceptives, different phases of the menstrual cycle) or reporting correlations with baseline hormone levels (Table 2). Cortisol responses were only examined across naturally occurring hormonal fluctuations (e.g., the menstrual cycle) and none of the studies involved an experimental manipulation of sex hormones. We found 12 studies that assessed progesterone (Antov & Stockhorst, 2014; Barel et al., 2018; Boisseau et al., 2013; Bürger et al., 2025; Childs et al., 2010; De Souza et al., 1991; Gaffey & Wirth, 2014; Galliven et al., 1997; Gordon & Girdler, 2014; Maki et al., 2015; Pletzer et al., 2021; Stephens et al., 2016) and 11 studies that assessed estradiol (Antov & Stockhorst, 2014; Barel et al., 2018; Boisseau et al., 2013; Bürger et al., 2025; De Souza et al., 1991; Galliven et al., 1997; Gordon & Girdler, 2014; Maki et al., 2015; Pico-Alfonso et al., 2007; Pletzer et al., 2021; Stephens et al., 2016), involving a total of 758 participants. Of the included studies, 3 compared naturally cycling women with women using oral contraceptives (Barel et al., 2018; Boisseau et al., 2013; Bürger et al., 2025). Five studies examined differences between women in the early follicular and luteal phases of the menstrual cycle (Childs et al., 2010; De Souza et al., 1991; Galliven et al., 1997; Gordon & Girdler, 2014; Maki et al., 2015), and 2 compared women in the early follicular phase with those in the mid-cycle/ovulatory phase (Antov & Stockhorst, 2014; Pico-Alfonso et al., 2007). Two studies compared men to women in the luteal phase (Barel et al., 2018; Pletzer et al., 2021), while one study compared men to women in the early follicular phase (Stephens et al., 2016). To avoid confounding the analysis by sex or gender differences that go beyond the role of the sex hormones, we only included correlations of the cortisol response with baseline progesterone or estradiol levels as effects of interest for each group separately in these studies (Barel et al., 2018; Pletzer et al., 2021; Stephens et al., 2016). All studies used a between-subjects design, except for one using a mixed within- and between-subjects design (Boisseau et al., 2013), while another one used a within-subject design (Gordon & Girdler, 2014). Most studies employed psychosocial stress tasks, 3 applied a physical stressor (Boisseau et al., 2013; De Souza et al., 1991; Galliven et al., 1997), and 1 used a combined psychosocial-physical stressor (Bürger et al., 2025).

**Table 2.** Characteristics of included studies investigating sex hormones and their effect on cortisol stress reactivity. *Note:* SD = standard deviation. NOS = Newcastle-Ottawa Scale for cohort studies. Study quality was categorized as low (0–4), moderate (5–7), or high (8–9). OC = oral contraceptives. IUD = Intrauterine Device.

| Author (year) | N | Cycle phase | Mean age (SD) | Percent women | Design | Hormone (saliva, blood) | Cortisol specimen | Stressor type | Time of cortisol measurement | Study quality (NOS) |
| --- | --- | --- | --- | --- | --- | --- | --- | --- | --- | --- |
| Antov et al. (2014) | 13 | early follicular phase | 23.9 (1.3) | 100 | between | estradiol (blood), progesterone (blood) | saliva | psychosocial | morning, afternoon | 7 |
| Barel et al. (2018) | 58 | luteal phase (n = 17), OC (n = 20) | 24.8 (2.5) | 63.8 | between | progesterone (saliva) | saliva | psychosocial | morning | 7 |
| Boisseau et al. (2013) | 16 | early follicular and mid-luteal phase (n = 7), OC (n = 9) | 22.8 (1.7) | 100 | between, within | estradiol (blood), progesterone (blood) | saliva | physical | morning | 5 |
| Childs et al. (2010) | 80 | follicular phase (n = 29), luteal phase (n = 23) | 21.9 (3.8) | 65.0 | between | progesterone (blood) | saliva, plasma | psychosocial | morning | 7 |
| Gaffey et al. (2014) | 38 | OC (n = 2), not mentioned (n = 18) | not provided | 52.6 | between | progesterone (saliva) | saliva | psychosocial | afternoon | 7 |
| Gordon et al. (2014) | 57 | early follicular, late follicular, luteal phases | 27.5 (0.8) | 100 | within | estradiol (blood), progesterone (blood) | plasma | psychosocial | morning | 7 |
| Maki et al. (2015) | 20 | early follicular phase | 25.6 (5.4) | 100 | between | estradiol (blood), progesterone (blood) | saliva | psychosocial | afternoon | 7 |
| Pico-Alfonso et al. (2007) | 18 | early follicular phase | 23.4 (2.4) | 100 | between | estradiol (blood) | plasma | psychosocial | morning, afternoon | 7 |
| Pletzer et al. (2021) | 67 | luteal phase | 22.9 (2.7) | 44.8 | between | progesterone, estradiol (saliva) | saliva | psychosocial | afternoon | 6 |
| Stephens et al. (2016) | 282 | follicular phase | 23.6 (3.3) | 47.9 | between | progesterone, estradiol (blood) | saliva & plasma | psychosocial | afternoon | 7 |
| De Souza et al. (1991) | 8 | early follicular, midluteal phase | 29 (4.24) | 100 | between | progesterone, estradiol (blood) | blood | physical | afternoon | 6 |
| Galliven et al. (1997) | 15 | follicular phase, midcycle phase, luteal phase | 30.1 (2.8) | 100 | between | progesterone, estradiol (blood) | blood | physical | morning, afternoon | 6 |
| Bürger et al. (2025) | 86 | IUD (*N* = 27), OC (*N* = 30), luteal phase (*N* = 29) | 23.6 (2.75) | 100 | between | progesterone, estradiol (blood) | blood, saliva | psychosocial-physical | afternoon | 8 |

### Coding procedure

Reviewer 1 (MK) screened the titles and/or abstracts of the studies identified through the search strategy to identify potential matches with the inclusion criteria. Any studies that could not be conclusively evaluated based on the title and/or abstract were marked for further assessment during the full paper review.

For each study, we extracted the reported cortisol levels and standard deviations before stress onset and the peak response and/or the area under the curve and standard deviation for all groups as outcome variables. In addition, we included correlation values with the baseline hormone levels if reported. We extracted relevant information on the study design: characteristics of participants (total number of participants, overall mean age, number of female participants, mean age of female participants, number of male participants, mean age of male participants), cortisol sampling method, the time of sampling relative to stressor onset, details regarding the experimental session (e.g., time of day) and the type of stressor, mean estradiol and progesterone, mean insulin, and blood glucose, as well as information on menstrual cycle phase and oral contraceptive usage (applicable to female participants only). If the data was not accessible from the article, the corresponding authors were contacted via email to request the necessary information. The data from all papers was extracted from at least two of the reviewers (MK, AK, and MM) and trained research assistants (RB, RL, SB, SG, and SB) independently, and inconsistencies were resolved by the third reviewer in discussion with the other reviewers to ensure accuracy and consistency.

### Risk of bias

To determine publication bias, a visual inspection of funnel plots (Sterne & Egger, 2001) and box plots was conducted. Assessments were done at the outcome level. Additionally, we used the open-access revised Cochrane risk of bias tool for randomized trials (RoB2, Sterne et al., 2019) to evaluate the metabolic studies in five different categories in low, medium and high risk. The categories consist of the randomization quality, the blinding, the effect of potential missing data, the validity of the method used to measure the outcome (e.g., if cortisol responses were objectively measured via biological assays), and whether the study followed a pre-registered analysis plan. Studies that were rated as high or medium risk of bias in more than one category were classified as having an overall high risk of bias. If only one category was rated as high or medium risk of bias, we classified the study as having a medium risk (Pervaz et al., 2025). In total (Fig. 2), 2 studies were rated as low risk of bias (Meier et al., 2021; Rüttgens & Wolf, 2022), 2 as medium risk (Bentele et al., 2021; Meier et al., 2025), and 4 as high risk of bias (Gonzalez-Bono et al., 2002; Kirschbaum et al., 1997; von Dawans et al., 2021; Zänkert et al., 2020). For the risk of bias assessment in studies on sex hormones (Table 2), we applied the Newcastle-Ottawa scale for cohort studies (NOS, Wells et al., 2014), considering menstrual cycle phase or oral contraceptive usage as the exposure variable, with participants subsequently assigned to stress or control tasks. NOS is a commonly used tool for evaluating the quality of non-randomized studies, which rates studies on three domains: selection of study groups, group comparability, and assessment of exposure or outcome, with a maximum possible score of 9. Based on the total score (Wang et al., 2024), study quality was categorized as low (0–4), moderate (5–7), or high (8–9) (Table 2).

**Fig. 2.**
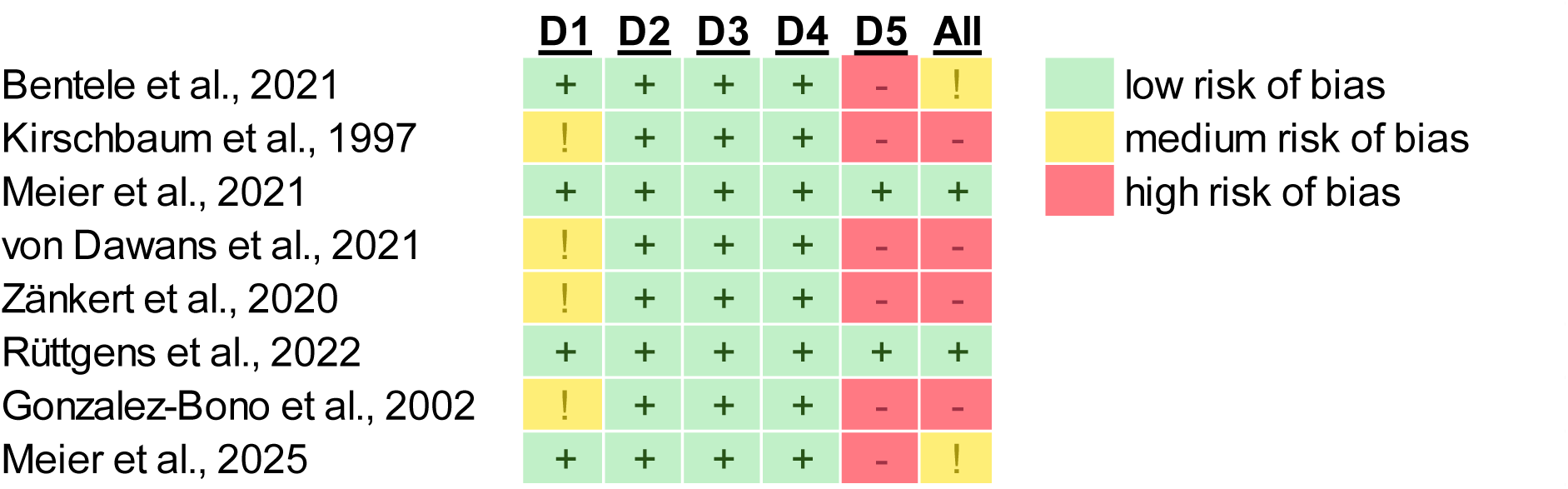
Risk of bias assessment of all included metabolic studies using a standardized checklist (Sterne et al., 2019). The studies were rated in five categories. D1: Randomization process. D2: Deviations from the intended interventions. D3: Missing outcome data. D4: Measurement of the outcome. D5: Preregistration.

### Statistical methods

All statistical analyses were performed using R (version 4.3.3). For the comparisons between glucose levels and cortisol responses or sex hormone levels and cortisol responses, we calculated effect sizes using Cohen’s *d* (Hedges & Olkin, 1984) based on means and standard deviation (*SD*) for all group comparisons or correlations with baseline cortisol levels (for non-manipulated sex hormones only) using the *metafor* package (Viechtbauer, 2010). If multiple effects were reported within one paper (e.g., numerous cycle phases were compared or peak cortisol response and area under the curve were reported), we aggregated all effect sizes with the same comparison of interest (low vs. high baseline hormone or glucose levels) within one paper using the Borenstein-Hedges-Higgins-Rothstein (BHHR, Borenstein et al., 2009) method, as implemented in the MAd package (Hoyt, 2014), assuming a correlation of 0.5 between effect sizes. Data manipulation was performed with the *dplyr* package (Wickham et al., 2023). Visualizations were created with *ggplot2* (Wickham, 2016) for outlier and Bayes factor robustness plots, and funnel plots were generated with the *metafor* package (Viechtbauer, 2010). Bayesian posterior distributions were visualized using half-eye plots with credible intervals generated with the *ggdist* package in R (Kay, 2025). Bayesian random-effects meta-analyses were conducted using the *bayesmeta* package (Röver, 2020), specifying a half-Cauchy prior (scale = 0.5) for the heterogeneity parameter (τ) and normal priors on the overall effect with *M* = 0 and *SD* = 0.75, 1.5, 2.5, and 3.5. Bayes factors for the null versus alternative hypotheses (BF_01_) were calculated to quantify evidence for the absence or presence of an effect, and robustness checks were performed across these prior choices. Outliers were identified using the interquartile range method (values beyond 1.5 × IQR from the first and third quartiles), and all analyses were repeated with and without outliers to evaluate the stability of results.

## Results

### Glucose intake increases stress reactivity

To assess the effects of glucose administration on cortisol stress reactivity, we compared groups that ingested glucose before a stress induction with control or placebo groups using Bayesian random-effects meta-analysis. As hypothesized, glucose intake was associated with a significant increase in cortisol stress reactivity across all studies including a glucose manipulation (*d* = 0.30, 95% CI = [0.05, 0.60], Fig. 3) with low to moderate heterogeneity among studies (*τ* = 0.14, 95% CI = [0.00, 0.52]), as further visualized in the posterior distributions (Fig. 7). This corresponds to a number needed to treat (NNT) of 10.6, meaning that glucose must be administered to approximately 11 individuals to observe one additional case of heightened cortisol reactivity compared to non-glucose conditions. The Bayes factor analysis provided anecdotal evidence for the effect of glucose on stress reactivity (BF_10_ = 2.42), and the robustness analysis showed that evidence was moderate at most under narrower priors and anecdotal with wider priors (Fig. 6). The funnel plot (Fig. 4) showed weak asymmetry. An outlier check identified 2 studies as potential outliers (Gonzalez-Bono et al., 2002; Kirschbaum et al., 1997). After excluding these studies, the meta-analysis still showed a small positive effect of glucose on cortisol stress reactivity (*d* = 0.22, 95% CI = [0.01, 0.45]), but the Bayes factor indicated anecdotal evidence against the alternative hypothesis (BF_10_ = 0.69), suggesting that additional evidence is needed to draw meaningful conclusions.

**Fig. 3.**
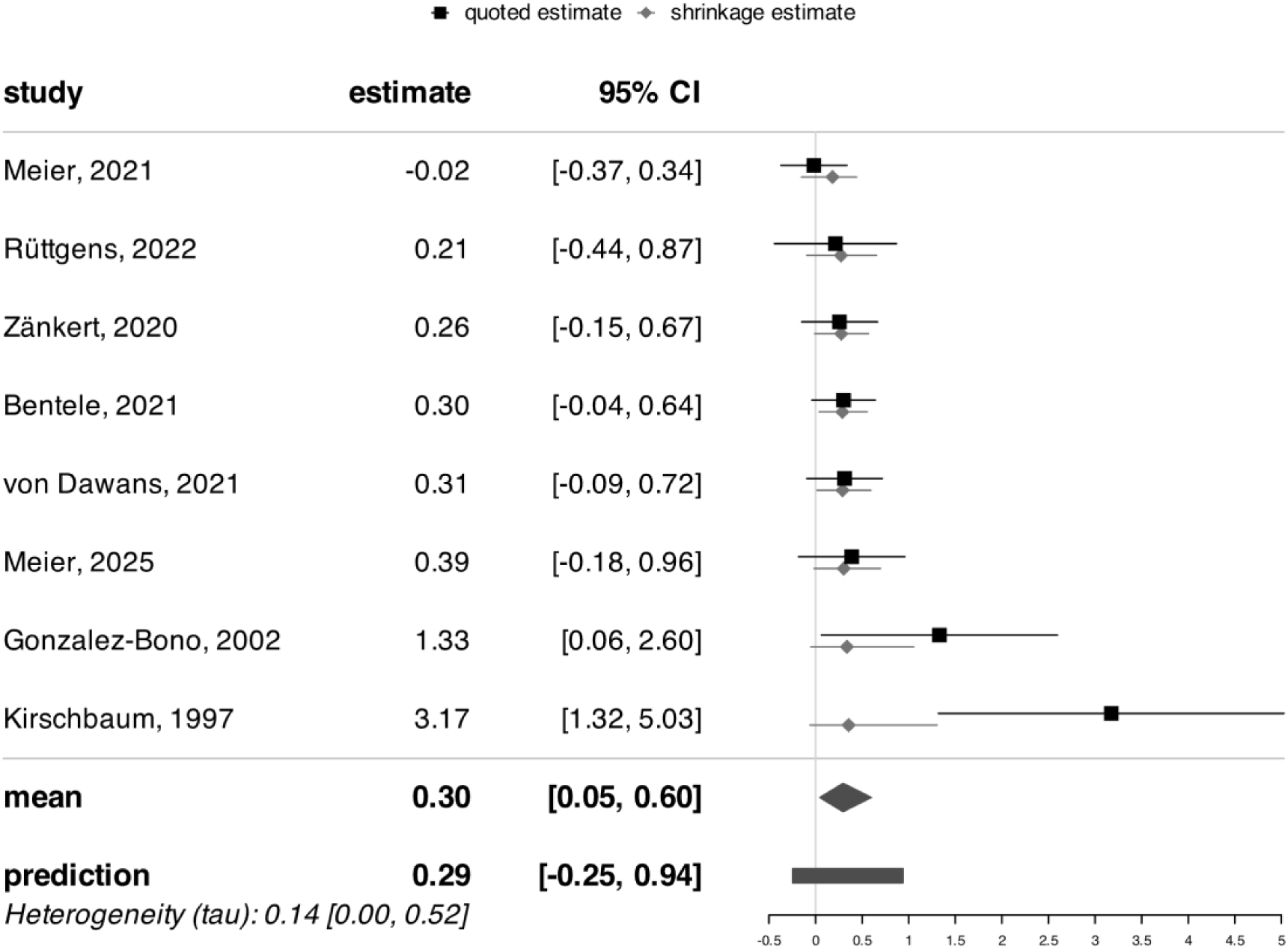
Forest plot of all included metabolic studies, showing an enhancing effect of glucose on cortisol stress reactivity (*d* = 0.30, CI = [0.05, 0.60], BF_10_ = 2.42) compared to control conditions (e.g., water). Individual study effect sizes (Cohen’s d) and 95% confidence intervals (CI) are displayed.

**Fig. 4.**
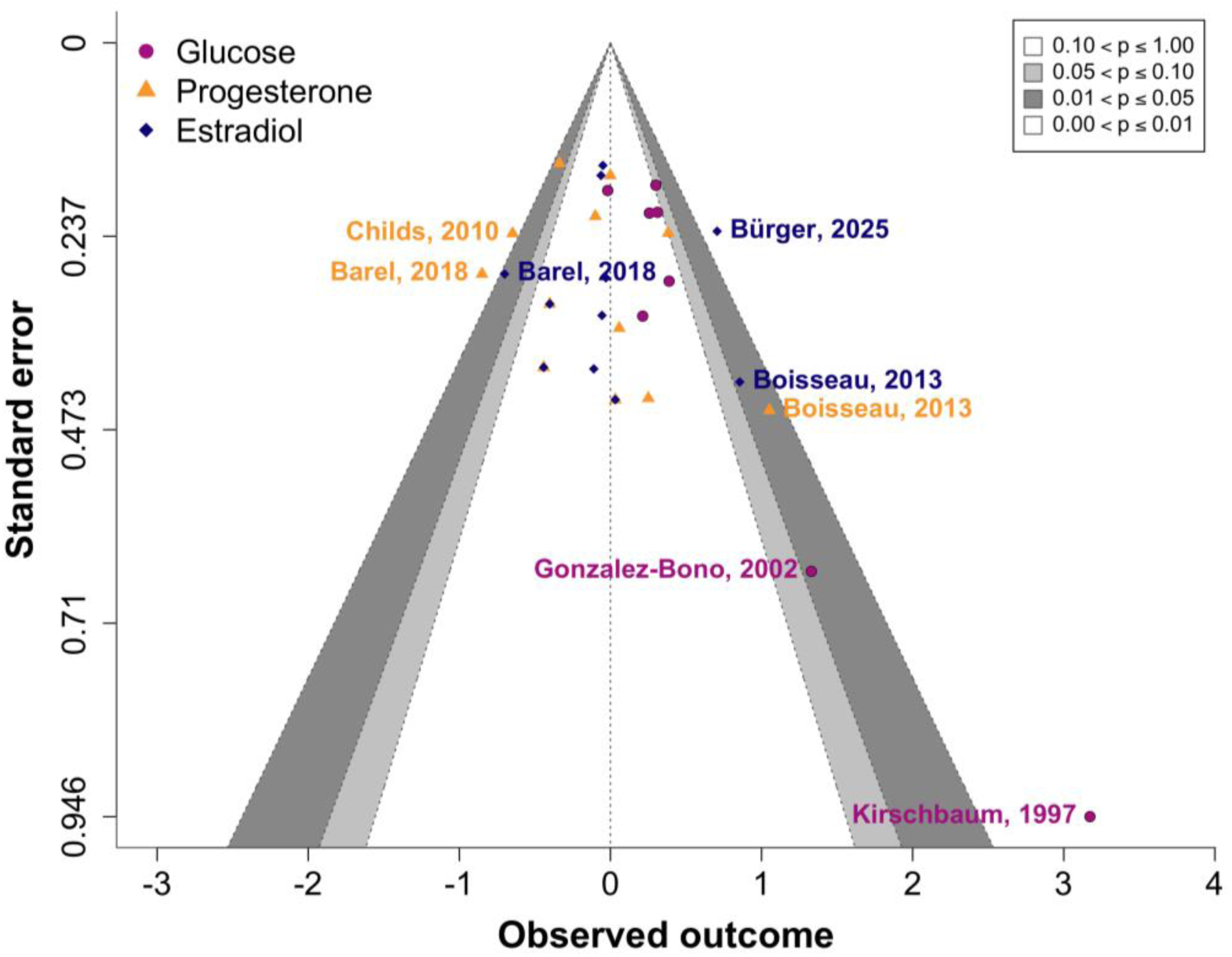
Contour-enhanced funnel plot showing standard errors and observed outcomes for each included study. Metabolic studies are depicted in magenta, studies assessing progesterone are shown in orange, and studies measuring estradiol are in blue.

**Fig. 5.**
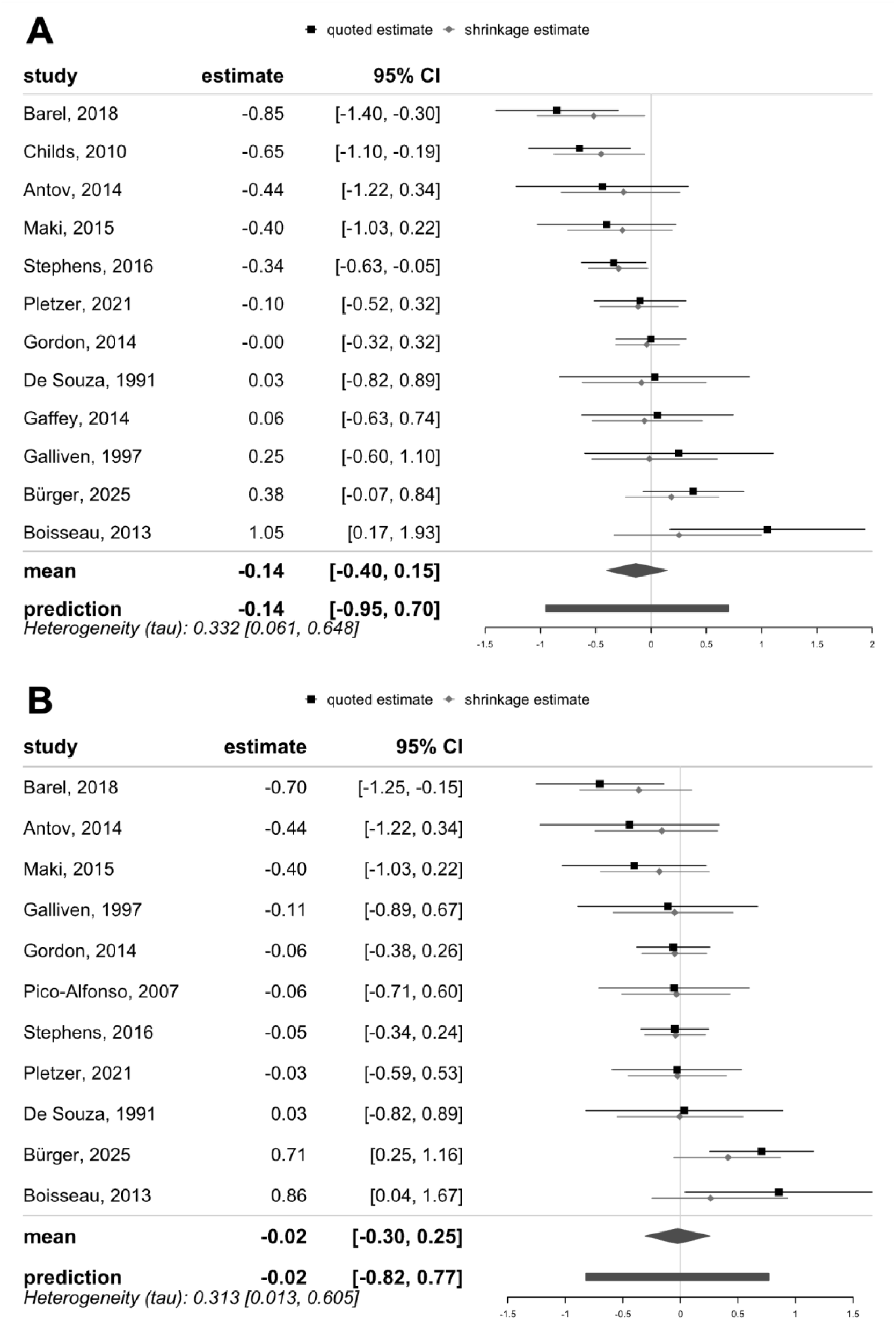
Forest plots of all included studies examining the relationship between sex hormone levels and cortisol stress reactivity. (A) Comparing high versus low progesterone levels, studies indicate no effect of higher versus lower progesterone levels on cortisol stress reactivity (*d* = -0.14, 95% CI = [-0.40, 0.15], BF_01_ = 6.01). (B) Assessing the effect of estradiol, studies show no significant difference in cortisol stress reactivity between higher and lower estradiol levels (*d* = -0.02, 95% CI = [-0.82, 0.77], BF_01_ = 11.64). Effect sizes are presented as Cohen’s d with 95% confidence intervals.

**Fig. 6.**
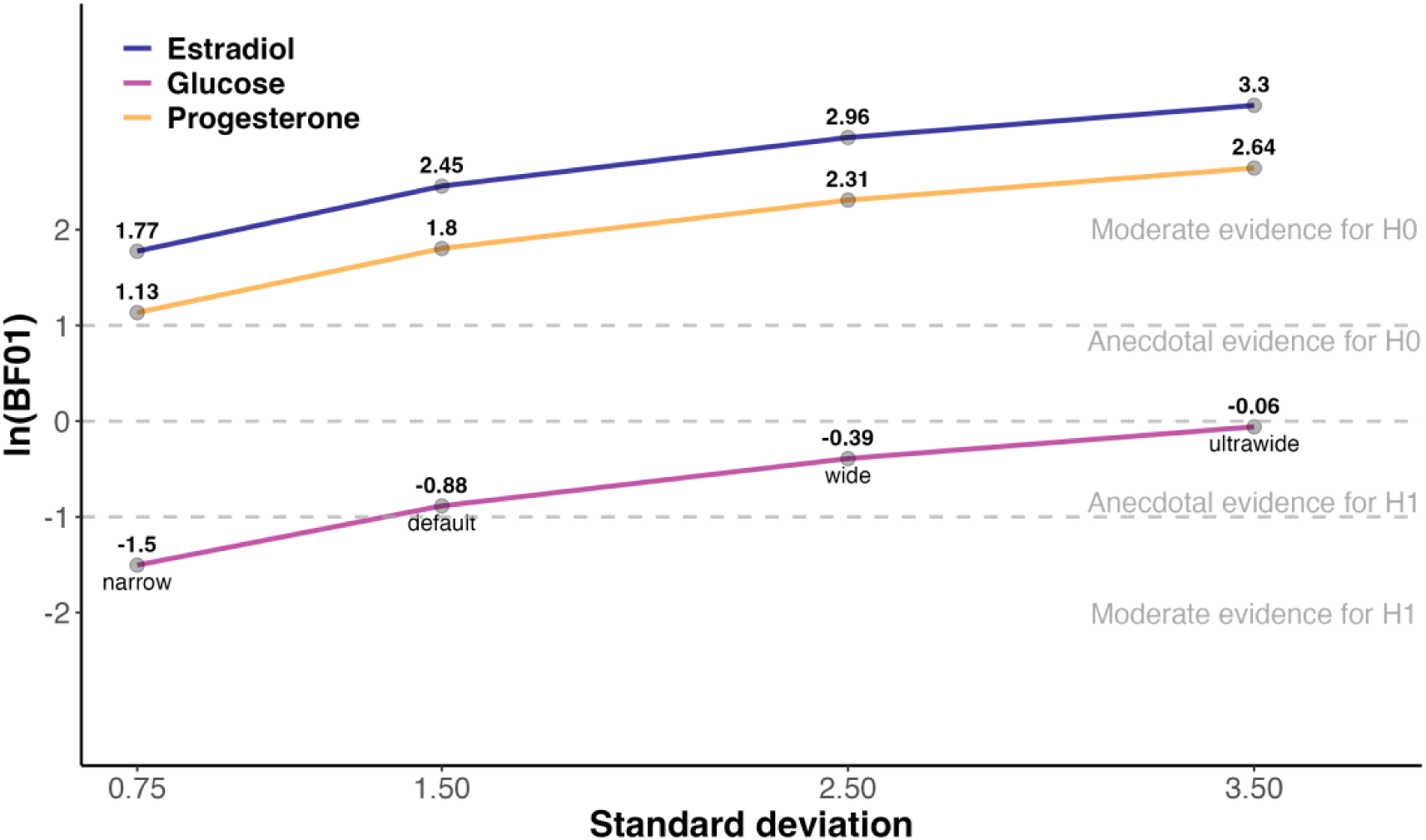
Bayes factor (BF_01_, log-transformed) robustness plot across different priors. The plot shows consistent moderate to strong evidence for H_0_ for estradiol (in blue) and progesterone (in orange) across all prior widths. For glucose (in magenta), evidence is inconsistent and prior-sensitive, shifting from moderate evidence for H_1_ with narrow priors to weak evidence for H_0_ with ultrawide priors.

**Fig. 7.**
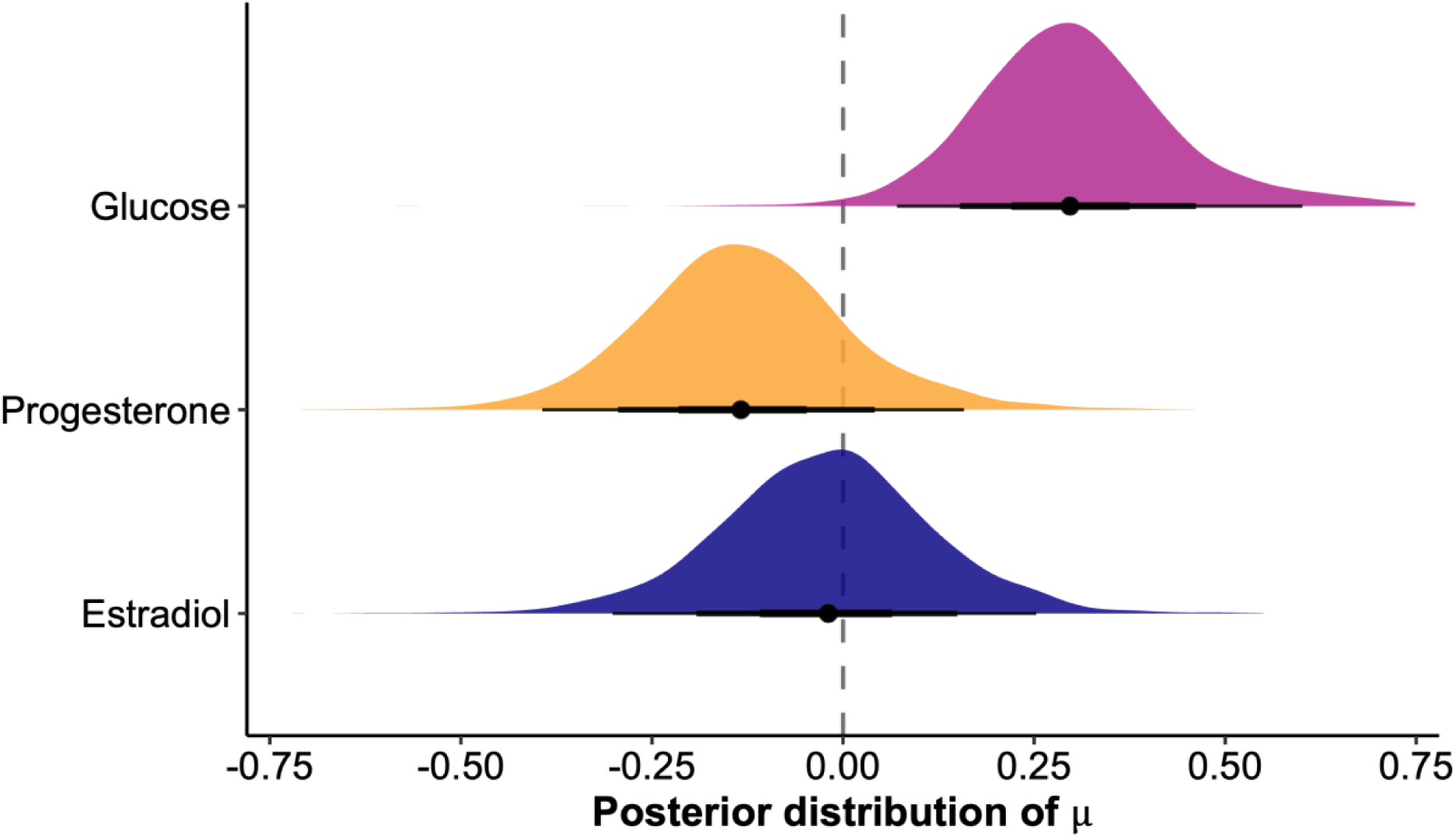
Bayesian posterior distributions of metabolic and hormonal effects on cortisol reactivity. The plot shows estimated posterior means and 95% credible intervals (black lines) for glucose (magenta), progesterone (orange), and estradiol (blue). Glucose was associated with a small positive effect on cortisol reactivity, while progesterone and estradiol showed no robust effects, with posterior distributions centered near zero and providing stronger support for the null hypothesis.

### Progesterone and estradiol levels do not robustly influence stress reactivity

To evaluate the role of sex hormone levels on stress reactivity, we performed two separate meta-analyses comparing groups with either low versus high progesterone or estradiol. First, higher progesterone levels were not significantly associated with cortisol stress reactivity (*d* = -0.14, 95% CI = [-0.40, 0.15], BF_01_ = 6.07, Fig. 5A), with moderate heterogeneity across studies (*τ* = 0.33, 95% CI = [0.06, 0.65]). The funnel plot suggested weak asymmetry (Fig. 4), indicating limited publication bias. One study (Boisseau et al., 2013) was identified as a potential outlier. When this study was excluded, the effect size remained non-significant (*d* = -0.20, 95% CI = [-0.44, 0.05]), and Bayesian analysis provided anecdotal evidence for the null hypothesis (BF_01_ = 2.78), which increased to moderate levels with wider priors. Similarly, estradiol levels showed no significant association with cortisol stress reactivity (*d* = -0.02, 95% CI = [-0.30, 0.25], BF_01_ = 11.64, Fig. 5B), with moderate heterogeneity (*τ* = 0.31, 95% CI = [0.01, 0.61]). The funnel plot showed no strong asymmetry (Fig. 4). Three studies were identified as potential outliers (Barel et al., 2018; Boisseau et al., 2013; Bürger et al., 2025). However, excluding these studies did not change the result (*d* = -0.10, 95% CI = [-0.31, 0.10], BF_01_ = 8.62).

## Discussion

Metabolic and hormonal states play a pivotal role in regulating adaptive stress responses in mechanistic studies, which may help explain individual variability in stress-induced human cortisol reactivity. To address this gap, we conducted a systematic review and meta-analysis examining differences in cortisol stress reactivity depending on baseline metabolic markers and sex hormones. Across all studies, we observed increases in stress-induced cortisol levels if glucose levels were higher (e.g., after a glucose load). In contrast, progesterone and estradiol were not consistently associated with stress-induced cortisol levels. Overall, our results highlight metabolic states as a robust modulator of cortisol reactivity, while associations with sex hormones appear less consistent to date.

As predicted, we found that glucose administration increased cortisol responses to acute stress. This aligns with studies showing that fasting tends to blunt HPA axis reactivity, whereas glucose intake restores cortisol reactivity (Bentele et al., 2021; Gonzalez-Bono et al., 2002; Kirschbaum et al., 1997). Crucially, studies comparing different macronutrients report larger cortisol increases following glucose or grape juice intake compared to fat or protein intake control conditions, indicating that the effects are likely driven by glucose (Bentele et al., 2021; Gonzalez-Bono et al., 2002; Zänkert et al., 2020). Preliminary evidence suggests that the type or intensity of the stressor may modulate this effect (Rüttgens & Wolf, 2022). However, individual changes in blood glucose levels after consumption do not correlate robustly with increases in cortisol, suggesting that circulating glucose alone may not fully explain the effect (Meier et al., 2021). Future studies should therefore not only focus on glucose-induced changes in blood levels but also consider individual variations in insulin sensitivity. Including measures of additional metabolic factors, such as insulin, triglycerides or ghrelin, may help to better evaluate individual differences in the potential to metabolize glucose efficiently. Taken together, our systematic review and meta-analysis suggest that glucose ingestion amplifies cortisol stress responses, but open questions remain concerning differences between various kinds of stress and individual differences in energy metabolism.

In contrast to the facilitating effect of glucose intake on stress-induced cortisol reactivity, the associations with sex hormones were less congruent across studies. Most included studies point to negative associations of progesterone and estradiol with cortisol reactivity. However, these effects did not reach significance in our analysis and provided only anecdotal evidence for the null hypothesis, indicating the need for more highly powered and conclusive studies. This contrasts with a prior meta-analysis reporting increased cortisol in women with high luteal-phase progesterone exposed to physical stressors (Klusmann et al., 2023), suggesting that hormonal effects may depend on the stressor type. Specifically, progesterone and cortisol seem to be co-released during physical stressors (Boisseau et al., 2013), while psychosocial stress paradigms often elicit reduced cortisol responses during high-progesterone phases (Maki et al., 2015; Stadler et al., 2019; Stephens et al., 2016; Wirth, 2011). Such differences between different types of stress likely reflect partially distinct neural pathways (Kogler et al., 2015). Estradiol findings were mixed, with no consistent evidence for a modulatory role. Inconsistencies across these studies may partly reflect variations in endogenous (e.g., menstrual cycle phase) and exogenous (e.g., hormonal contraceptive use) hormonal modulation across studies (Boisseau et al., 2013), which were often not contrasted. Other potential modulators, such as testosterone, and factors influencing hormone bioavailability, including sex hormone-binding globulin and corticosteroid-binding globulin, may also influence cortisol reactivity (Bürger et al., 2025). Overall, the reviewed set of findings highlights the current ambiguity of findings, calling for improved study designs to conclusively confirm or refute the role of sex hormones in shaping stress-induced cortisol reactivity.

While this meta-analysis provides a quantitative synthesis of existing literature, some limitations should be considered. Methodological variability across studies, particularly regarding cortisol sampling timing, likely contributed to heightened heterogeneity in effect sizes. While most metabolic studies collected samples in the afternoon or evening to avoid confounding with the cortisol awakening response, sex hormone studies varied more, with 6 studies assessing cortisol in the morning (Barel et al., 2018; Boisseau et al., 2013; Childs et al., 2010; Galliven et al., 1997; Gordon & Girdler, 2014; Pico-Alfonso et al., 2007). This may have led to an overlap with the cortisol awakening response, with potential effects on stress response magnitude (Stalder et al., 2016). Additional variability may have stemmed from inconsistent menstrual cycle classifications and hormonal contraceptive use. Few studies systematically distinguished between these factors, often omitting contraceptive use type or duration and relying solely on self-reported cycle phase without hormonal verification, which undermines comparability across studies, as noted in previous meta-analyses (Hamidovic et al., 2020; Klusmann et al., 2023). Such inconsistencies persist despite findings that naturally cycling women and oral contraceptive users differ in cortisol responses to psychosocial stress (Gervasio et al., 2022). Moreover, although most metabolic studies included experimental manipulations and control conditions, sex hormone studies were primarily correlational. Several studies had small sample sizes, increasing the likelihood of sampling error and inflated effect size estimates (Stone & Rosopa, 2017). Future studies should address these limitations by using within-subject designs and, if feasible, experimentally manipulating hormone levels.

In addition to these methodological concerns, a notable conceptual gap is the lack of studies examining the interaction between metabolic state and sex hormone levels in the context of the acute stress response. No available study has systematically investigated whether the effects of sex hormones on cortisol stress reactivity differ depending on metabolic conditions such as fasting, glucose, or insulin sensitivity. Although one study measured both glucose and progesterone across different menstrual cycle phases (Galliven et al., 1997), they focused on glucose responses to physical stress rather than their combined effect on cortisol stress reactivity. Recent research suggests that glucose and insulin sensitivity fluctuate across the menstrual cycle (Hummel et al., 2023; Kroemer, 2023; Lin et al., 2023), pointing to a dynamic interaction between sex hormones and metabolic function. This may be relevant for the HPA axis regulation since changes in stress reactivity with increasing body mass index (BMI) are predominantly observed in females (Kühnel et al., 2023). This interplay may be crucial for understanding sex differences in stress-related disorders, particularly in populations with combined metabolic and hormonal dysregulation, such as individuals with obesity, metabolic syndrome, or type 2 diabetes, where stress responses and disease risk differ by sex (Navarro et al., 2015; Pradhan, 2014; Tronieri et al., 2017). Our findings showing more consistent effects of glucose than of sex hormones on cortisol suggest that metabolic signals may play a stronger or more direct modulatory role than sex hormones.

In conclusion, this meta-analysis highlights a robust enhancing effect of glucose on cortisol stress reactivity, while associations with sex hormones remain too inconsistent across studies, possibly due to heterogeneity in methods and the predominant reliance on observational studies. These findings emphasize the need for future interventional studies to systematically examine how metabolic and hormonal systems interact in modulating HPA axis responses. Understanding this interplay may offer important insights into sex differences in stress-related disorders and guide improved treatment approaches that incorporate state-dependent effects.

## Acknowledgments

This work was performed as part of the International Research Training Group: Women’s Mental Health Across the Reproductive Years (IRTG 2804). The authors thank Renée Boehm, Ricarda Lepsius, Selina Braun, Svea Gründel, and Sandra Beinbauer for their assistance in extracting the data. MK was supported by the International Research Training Group: Women’s Mental Health Across the Reproductive Years (IRTG 2804) funded by the German Research Foundation (DFG); grant number: GRK 2804/1. Additionally, the meta-analysis was supported by the DFG grant KR 4555/9-1.

## Author contributions

NBK was responsible for the study concept and design. MK, MM & AK collected data under supervision by NBK. AK & NBK conceived the method and AK processed the data. AK & MK performed the data analysis and NBK contributed to analyses. MK, AK & NBK wrote the manuscript. All authors contributed to the interpretation of findings, provided critical revision of the manuscript for important intellectual content and approved the final version for publication.

