## supporting information for "Metabolic state shapes cortisol reactivity to acute stress: A systematic review and meta-analysis of metabolic and hormonal modulators"

**Supplementary Material**

*Inclusion and exclusion criteria*

Experimental studies were included in the analysis if they met the following criteria: (1) for cortisol stress reactivity, subjects must have been exposed to an acute stress induction using a standardized stress task or stressor; (2) measurements of cortisol levels should be done before and after stressor exposure (3) measurement of metabolic or sex hormone levels or groups with different sex hormone levels (e.g., in different menstrual cycle phases) were compared; (4) participants were 18 years and older, male or female; (5) at least one of the following parameters was measured or manipulated: estradiol, progesterone, insulin, glucose; (6) if measuring insulin, glucose, estradiol or progesterone, the assessment should be conducted before exposure to the stressor; (7) cortisol was measured in saliva or blood (serum and plasma).

Studies were excluded when one of the following criteria was met: (1) measurement of general or chronic stress levels, not situational stress; (2) the cortisol awakening response or oxidative stress was measured; (3) participants were younger than 18 years; (4) non-healthy sample (e.g., any mental, neurodevelopmental, endocrinological, or neurological sample); (5) non-English or German-published studies; (6) articles were non-original (e.g., reviews, meta-analyses, commentaries and editorials); (7) full text of the article was not available online, nor upon request from the corresponding authors.
